# ConVarT: a search engine for matching human genetic variants with variants from non-human species

**DOI:** 10.1101/2021.01.07.424951

**Authors:** Mustafa S. Pir, Halil I. Bilgin, Ahmet Sayici, Fatih Coşkun, Furkan M. Torun, Pei Zhao, Yahong Kang, Sebiha Cevik, Oktay I. Kaplan

## Abstract

The availability of genetic variants, together with phenotypic annotations from model organisms, facilitates comparing these variants with equivalent variants in humans. However, existing databases and search tools do not make it easy to scan for equivalent variants, namely “matching variants” (MatchVars) between humans and other organisms. Therefore, we developed an integrated search engine called ConVarT (http://www.convart.org/) for matching variants between humans, mice, and *C. elegans*. ConVarT incorporates annotations (including phenotypic and pathogenic) into variants, and these previously unexploited phenotypic MatchVars from mice and *C. elegans* can give clues about the functional consequence of human genetic variants. Our analysis shows that many phenotypic variants in different genes from mice and *C. elegans*, so far, have no counterparts in humans, and thus, can be useful resources when evaluating a relationship between a new human mutation and a disease.

## INTRODUCTION

The progress of the genomics field is accelerating with the aid of next-generation sequencing technologies, most recently exemplified by the launch of the Exome Aggregation Consortium (ExAC) and Genome Aggregation Database (gnomAD). These genomic databases were gathered from more than 140,000 randomly selected individuals, providing large-scale collections of human genetic variants (1, 2). Humans possess over 700 million genetic variants in the protein-encoding and non-protein-coding regions of the genome. However, the growing number of human genetic variants does not automatically mean that our understanding of the functional effects of these variants concurrently increases (3). Due to their clinical importance, functional consequences are only available for a handful of genetic variants, while biological impacts are unavailable for most human genetic variants (4). The biological understanding of these genetic variants is crucial to diagnose diseases better and make therapeutic decisions regarding diseases, particularly noting each variant’s especially discerning contribution to rare and complex diseases.

Though examining the functional effects of variants is an efficient strategy for categorizing them as being either pathological or benign, a considerable workload and financial demand are limiting factors for such experimental efforts (5–9). In this regard, model organisms are regularly used to examine the functional influence of clinically relevant variants, but unprecedented quantities of variant and phenotype data from model organisms largely go unnoticed. We assume that when an amino acid, where it shows a similarity with other homologous sequences, undergoes an intolerable change in non-human species, as observed with phenotype-changing variants, human-equivalent variants will likely disrupt the function of genes in humans. The basis for this claim is that the function of proteins is likely to be compromised when the positions of highly conserved amino acids are altered (10). Despite various orthology available search, there is no existing search engine that: 1) allows users to scan for equivalent variants between human and non-human species; 2) incorporates available experimental data from model organisms into corresponding variant positions; and 3) enables users to submit the empirical variant data (11–15).

Here, we develop an integrated search tool and database called ConVarT (**Con**gruent clinical **Var**iation Visualization **T**ool) to search for equivalent variants (pathological variants, phenotypic variants, and variants of uncertain significance) between human and non-human species. As a proof of concept, we matched amino acid-changing variants from humans with variants from mice and the nematode *Caenorhabditis elegans* and generated a list of equivalent variants called matching variants (MatchVar) between human and mice or *C. elegans*. Furthermore, we integrated available annotations (pathological, phenotypic, etc.) for variants into the database, typically making it easy for users to search for MatchVars with phenotypic data. Through the use of MatchVars associated with functional data, our comprehensive database can provide an additional layer of proof for the functional interpretation of human variants.

## RESULTS

### ConVarT: a search tool for “matching variants”

The availability of organism-specific databases (WormBase for *C. elegans* and Mutagenetix for mice) containing non-naturally occurring variants and phenotype data for *C. elegans* and mice enables us to equate them with human genetic variants from gnomAD, Clinical Variants (ClinVar), COSMIC, and dbSNP **(Figure 1)** (2, 16–20). We initially derived the human gene orthologs for *M. musculus* and *C. elegans*. We then collected amino acid-changing variants from gnomAD, ClinVar, COSMIC, and dbSNP for humans, from WormBase for *C. elegans*, and from the Australian Phenome Bank and Mutagenetix for mice. gnomAD describes over 16,179,380 variations, while the ClinVar database deposits over 500,000 variants, including pathogenic and benign variants (2, 20). COSMIC and dbSNP contain over 6,842,627 variations and 1,086,546 variations, respectively (16, 19). Our analysis revealed that gnomAD, ClinVar, COSMIC, and dbSNP have identical and non-identical variants (**Figure 2A)**. Mutagenetix and WormBase present almost 800,000 variants from these organisms (17, 18). To compare human variants with those from mice and *C. elegans*, we matched the protein sequences of the ortholog genes, followed by performing 2,379,397 multiple sequence alignments (MSAs) of all combinations of these protein sequences to ensure alignment of the corresponding amino acid positions. The schematic workflow of ConVarT is presented in **Figure 1.** We then integrated genetic variants and variant-specific annotations (such as pathogenicity, phenotypes, and allele frequency) from humans, mice, and *C. elegans* into corresponding amino acid positions in each gene. We restricted our comparisons to amino acid-changing variants, as amino acids are more conserved than non-amino acid-changing variants, and protein function is likely to be affected by an alteration in the corresponding amino acids. For visualization of MSAs together with variants from humans, mice, and *C. elegans*, we used in-house software (21). Previous studies revealed that many human variants overlap with amino acids undergoing post-translational modifications (PTMs) and are enriched at the protein domains. Therefore, Pfam protein domains for humans and 383k PTMs from PhosphoSitePlus were systematically integrated into corresponding positions (22, 23).

**Figure 1:**
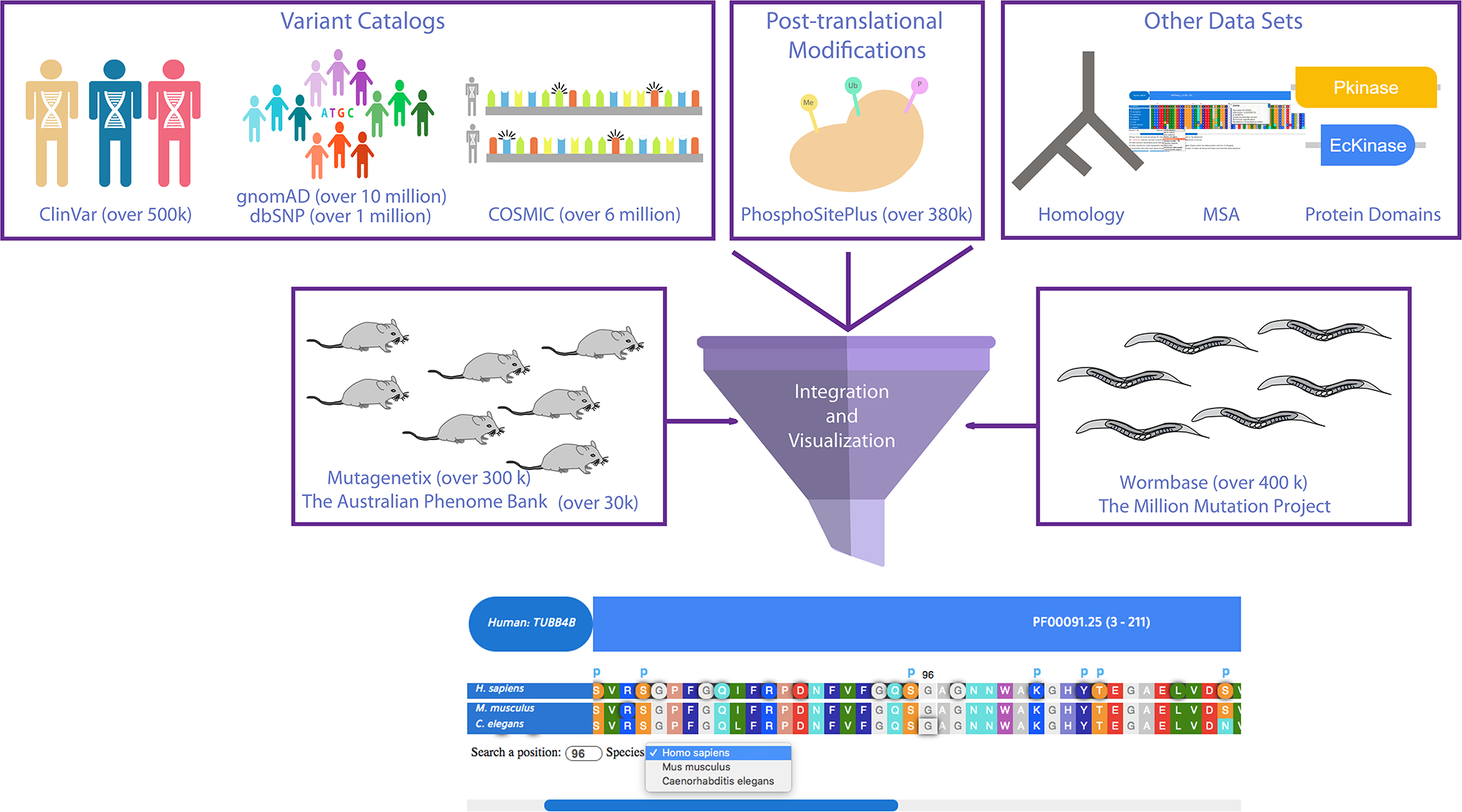
The schematic flow of ConVarT, a search engine for matching variants. Gene orthology was determined, followed by MSAs and the incorporation of human variants from gnomAD, ClinVar, COSMIC and dbSNP, and variants from the Australian Phenome Bank (Mice), Mutagenetix (Mice) and WormBase (*C. elegans*). Human PTMs and human protein domains are integrated for co-visualization with the MSA and variants.

**Figure 2:**
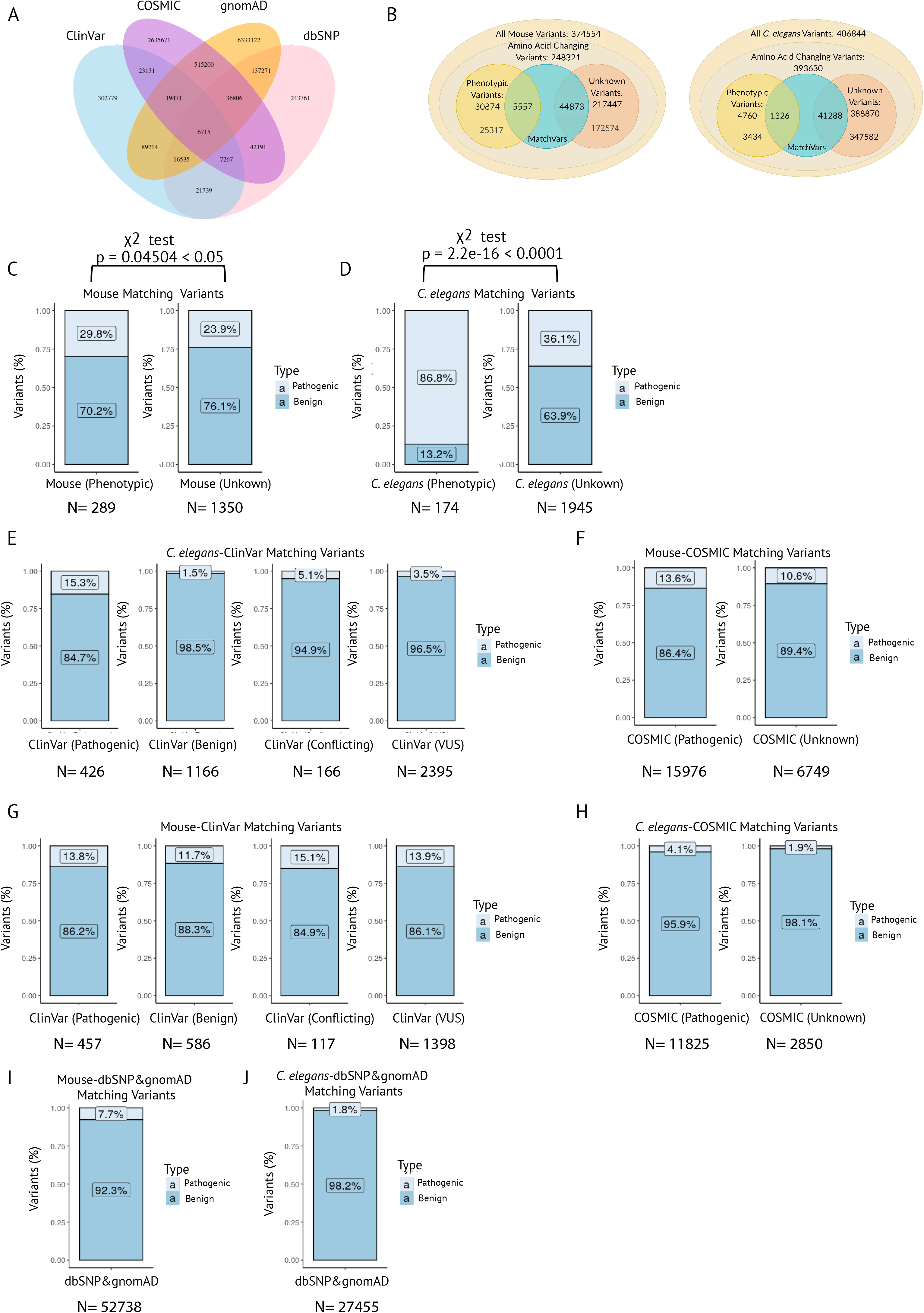
Matching variants among human, mice, and *C. elegans*. **A)** Venn diagram showing the number of overlapping and non-overlapping variants between gnomAD, ClinVar, COSMIC, and dbSNP **B)** the total number of MatchVars with phenotypes and unknown significance from mice and *C. elegans* were displayed in the two distinct Venn diagrams. **C), D),** Bar plots display the fraction of phenotypic or unknown significance of MatchVars from mouse and *C. elegans* with either ClinVar pathogenic or benign variants. A statistically significant difference between two proportions was displayed for mouse and *C. elegans.* X^2^= Chi Square **E), F) G), H, I and J)** Bar plots display comparisons of phenotypic and unknown MatchVars from mice and *C. elegans* (percentage) with a range of human variants databases ClinVar, COSMIC, gnomAD, and dbSNP. N: number of MatchVars between humans, mice or *C. elegans*.

Our analysis revealed that many equivalent amino acid substitutions exist between humans and these two organisms (**Figure 2B**). For example, both human AP1M1 and *C. elegans* UNC-101, the homologue of AP1M1, have the identical residue change at the corresponding positions (Human NP_001123996.1: p.P375S, Variant ID: rs1490425955 vs *C. elegans* NP_001040675: p.P362S, Variant ID: WBVar00679460). We have named these equivalent variants “matching variants” (MatchVars) (**Figure 2B, and Table S1, S2**). For a variant to be considered a MatchVar, the amino acid residue at the corresponding positions must be matched between the human gene and the ortholog genes. The Materials and Methods provide a detailed explanation. There are 50430 MatchVars between humans and mice, while the number of MatchVars between humans and *C. elegans* is 42614 (**Figure 2B, and Table S1, S2**). Finally, to make the MatchVars available to the community, we created a web-based search engine and database called ConVarT (http://www.convart.org/) to facilitate consistent and rapid visualization of MatchVars and variant annotations on MSAs together with human PTMs and protein domains. ConVarT offers a protein sequence similarity index based on the matching residues for three species below part of MSAs so that the researchers utilize this section to gain more insight into the conservation of the sequence in the model organism compared to humans.

ConVarT reduces the time needed to find the phenotypic and unknown significance of MatchVars in the genomes of three organisms from hours to seconds. A comprehensive list of identifiers that can be used to search for genes on ConVarT can be found in **Table S17**. Furthermore, users can now submit any protein sequence as an input, and ConVarT will perform sequence comparison and alignment to find the closest human orthologue of that protein sequence, as well as display human variants on the pairwise sequence alignment of human protein and submitted protein sequence (**Figure S2)**. This is especially useful for model organism researchers, such as *S. cerevisiae,* who have yet to be incorporated into ConVarT.

### Usage of ConVarT to conjecture about the functional effects of variants in disease analysis

Disease-causing variants are annotated as pathogenic in humans, whereas variants with phenotypic annotations in non-human species indicate disease-like conditions for them, suggesting that both types of annotations represent the disruption of the protein function. The majority of human genetic variants in the ClinVar database are single nucleotide variants (SNVs), and variant of uncertain significance (VUS) make up the majority of these SNVs (**Figure S1A and B)**. Expectedly, compared to the fraction of phenotypic MatchVars coinciding with benign variants of ClinVar, the proportion of phenotypic MatchVars of *C. elegans* overlapping with pathogenic variants of ClinVar is calculated to be significantly higher (p < 0.0001; Chi-square X^2^ test), which is consistent with the equality of pathogenic with phenotypic variants. For phenotypic MatchVars from mice, the mouse phenotypic MatchVars/ClinVar pathogenic variants ratio is slightly higher than the ratio of the mouse phenotypic MatchVars/ClinVar benign variants (p < 0.05; X^2^ test) (**Figure 2C and 2D**). ConVarT can serve researchers and human geneticists in several ways. First, they can assess whether variants of interest from humans currently have MatchVars already associated with the phenotype in the mice or *C. elegans,* potentially adding a new layer of support to their study. For example, the MatchVar p.G368E (allele name n4435 and NP_495455) in *C. elegans* KAT-1 (Human ACAT1) was identified as phenotypic (short life span) in 2010. The p.G388E variant (NP 000010) in human ACAT1 was pathogenic seven years later, resulting in human acetyl-CoA acetyltransferase deficiency (24, 25). We presented all the phenotypic MatchVars of mice and *C. elegans*, overlapping with pathogenic and other variants in ClinVar, COSMIC, dbSNP, and gnomAD (**Figure 2E, F, G, H, I and J, and Table S1- S16**). Second, there may currently be no MatchVars from humans for the phenotypic MatchVars of the mice or *C. elegans*. Where there may be the appearance of a new human variant associated with a disease, scientists and human geneticists may look for the phenotypic MatchVars in ConVarT, thereby helping to assign certain variants as being potentially detrimental. Finally, the Million Mutation Project and Mutagenetix have many strain collections that bear unique MatchVar mutations. Many of these mutations are poorly characterized, such as *C. elegans* and mice mutants bearing MatchVars (*C. elegans* DYF-18/CDK7 NP_502232: p.P245L and p.R26K; *C. elegans* OSM-3/KIF17 NP_001023308: p.T89I, p.P161L, p.S208L, p.A464T, and p.W572*; Mice Ros1, NP_025412: p.T220A and p.V2050A; Mice Tg, NP_033401: p.S164G, p.T1555S, and p.C1993R) (**Figure 3A, B, C, and D**) (17, 26). Therefore, the functional significance of these MatchVars is currently unknown. These mutants can be obtained from the distribution centers for functional analysis (17, 26). Researchers can also easily submit their variants and phenotype data to ConVarT, thereby sharing their phenotypic and variant data with the community.

**Figure 3:**
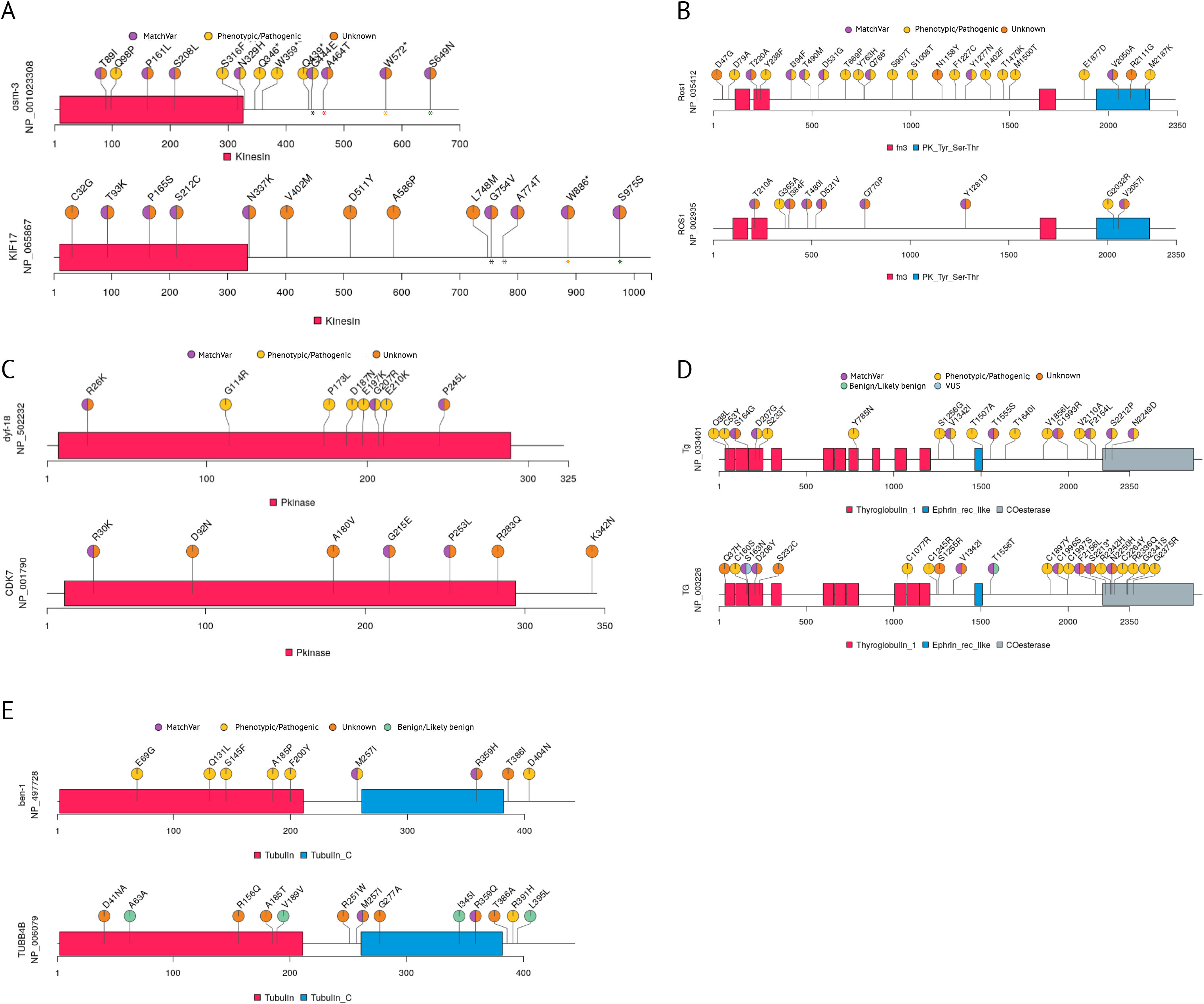
Distribution of matching and pathological/phenotypic variants per gene. Graphical representations of data (lollipop plots) exhibits pathological/phenotypic, orthologous, variants of unknown significance, and benign or likely benign variants (annotated amino acid changes) at their corresponding positions per gene. The length of each protein is shown, together with specific functional protein domains (different colors). Protein IDs and gene names are displayed next to the plots. **A), C), and E)** Comparison of *C. elegans* genes with human counterparts. In Figure 3A, different colors for the asterisk reflect MatchVars among humans and *C. elegans*. **B) and D)** Mice and human genes are displayed.

### Utilizing phenotypic variants from non-human species to add one layer support for understanding human variants

Eight missense variants in *C. elegans ben-1* (Human *TUBB4B*) encoding tubulin beta chain (NP_497728: p.E69G, p.Q131L, p.S145F, p.F167Y, p.A185P, p.F200Y, p.M257I, and p.D404N) were implicated in displaying resistance to benzimidazole (BZ) and were labeled as putative loss-of-function (LoF) variants (27, 28) (**Figure 3E**). We found that *C. elegans* missense residues were conserved in humans and mice, except for p.F200. Next, we investigated whether these conserved residues had MatchVars in humans, which revealed no MatchVars for p.E69G, p.Q131L, p.S145F, p.A185P, and p.D404N in humans. This result suggested that these variants were under negative selection in the human population, potentially due to their detrimental effects. All these variants were consistently predicted to be deleterious by SIFT (**Table S3)**. However, p.M257I and p.F167Y in *C. elegans* ben-1 (NP_497728) are the MatchVars of p.M257I and p.F167Y (variant IDs: COSM3929966 and COSM6451925) in human TUBB4B (NP_006079.1), respectively, predicted to be pathogenic by the FATHMM (**Figure 3E)** (19). Indeed, the absence of the p.M257I and p.F167Y mutations in human TUBB4B in gnomAD and TOPMed suggests that they were likely negatively selected. This data indicates that the experimental data already available from *C. elegans* and mice may provide additional evidence to deduce the functional implication of human variants together with allele frequency.

## DISCUSSION

Model organisms have contributed significantly to our knowledge of human biology and diseases. For this reason, the Alliance of Genome Resources recently launched an integrated database that shares genetic and functional genomic data from a variety of model organisms in order to aid scientists in their understanding of human biology and disorders, though they did not provide a search engine for MatchVars (29). This current work fills this gap by presenting a search engine for MatchVars and a database comparably displaying the pathogenic or phenotypic and unknown significance of variants from humans, mice, and *C. elegans.* ConVarT was designed with the goal of expanding and incorporating variants from other organisms, such as Zebrafish and Drosophila. ConVarT recently added 24,658,526 human variants from TOPMed to its database (**Table S18)** (3). The model organism community can easily submit new variant data and expand ConVarT to become an open platform for exchanging information about new MatchVars, thereby aiding the presentation of empirical evidence from model organisms to provide one more layer of support for interpretation of human variants.

## Material and Methods

### The gene homology list for humans and non-human species

The gene homology list for human and non-human organisms was first build using organism-specific orthology resources such as DIOPT, MGI, and the Zebrafish Information Network (ZFIN) (20, 31,33). In addition, as we uncovered inaccuracies and missing data in the orthology data, we manually corrected them using a reciprocal BLAST (35) analysis to find the counterparts of human genes. For example, the reciprocal BLAST analysis reveals that *C. elegans F42G8.19 (ceph-41)* is the most likely ortholog of human CEP41, *C. elegans* F42G8.19 does not emerge as an ortholog of human CEP41 based on orthology searches from *C. elegans* gene orthology database (32, 34).

### Download of Genetic Variants for Humans, Mice and *C. elegans* and Posttranslational modification (PTMs)

The human variants and organisms specific variants were downloaded from the following resources: ftp://ftp.ncbi.nlm.nih.gov/pub/clinvar/ (ClinVar), http://gnomad.broadinstitute.org/ (Aggregation Database (gnomAD v2.03), https://cancer.sanger.ac.uk/download (COSMIC), https://ftp.ncbi.nih.gov/snp/ (dbSNP) and https://www.phosphosite.org/Supplemental_Files (PhosphoSitePlus) (2, 16, 19, 20, 23). Mutagenetix (https://mutagenetix.utsouthwestern.edu/about.cfm) and Wormbase generously shared variants and phenotypic data for Mouse and *C. elegans,* respectively (17–18). We obtained mice phenotypic data and variants from the Australian Phenomics website (the webpage for download is https://pb.apf.edu.au/phenbank/homePage.html).

### Multiple Sequence Alignments

We retrieved human protein sequences and human homolog genes from chimp, Macaca, rat, mouse, zebrafish, *C. elegans, D. melanogaster,* and frog, and used ClustalW v2.1 pairwise alignment to perform 2,379,397 distinct combinations of multiple sequence alignments (MSAs) of these protein sequences (36). The final files were stored in MySQL together with variants from humans and non-human species.

### Data Visualization and Website Development

The recently developed MSA visualization tool was used to co-visualize MSAs along with annotations (variants and phenotypic data) (21). We constructed the ConVarT website and deposited all data for the users. Furthermore, we shared all of the necessary code (ConVarT Web https://github.com/thekaplanlab/ConVarT_Web) on GitHub.

Additional materials and methods are available in the supplementary file.

## Code availability

All codes used for generating data, websites, and figures are accessible from https://github.com/thekaplanlab.

## ACKNOWLEDGEMENT

We thank Dr. Seref Gul for critical reading of manuscripts. We gratefully thank Mutagenetix and Wormbase for generously sharing variants from Mutagenetix and Wormbase.

## AUTHOR CONTRIBUTIONS

O.I.K. and S.C originated project

H.I.B., F.M.T., and M.S.P. generated MSA data and collected variants

F.M.T. generated the homology table

H.B., F.T., A.S. and F.C. designed and developed web application

M.S.P. analyzed the data and generated Figure 2 and 3 and supplemental tables and figures

P.Z and Y. K. provided reagents.

O.I.K. directed the work of H.B., F.M.T., M.S.P., A.S. and F.C. and wrote the original draft

## FOOTNOTES

The authors declare no conflict of interest.

