## Supplementary Files for "ConVarT: a search engine for matching human genetic variants with variants from non-human species"

### Supplementary Figures

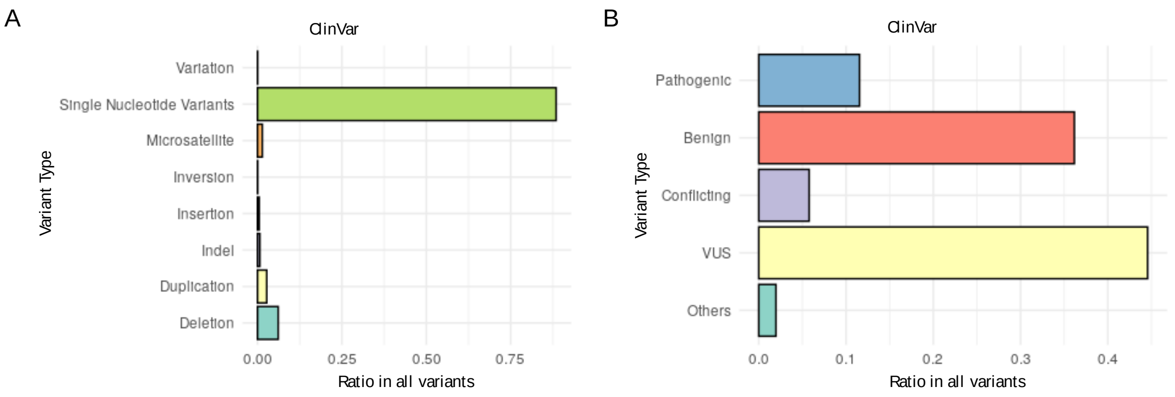


**Figure S1: A)** Distribution percentage of human genetic variants (792,968 variants) in the ClinVar database (downloaded on January 3, 2021). Single nucleotide variants (SNVs) make up the majority of the human genetic variants in the ClinVar database, followed by deletions. **B)** Graph displays the percentage breakdown of SNPs (701,905 variants) in the ClinVar database, including pathogenic, benign, and variants of uncertain significance (VUS).

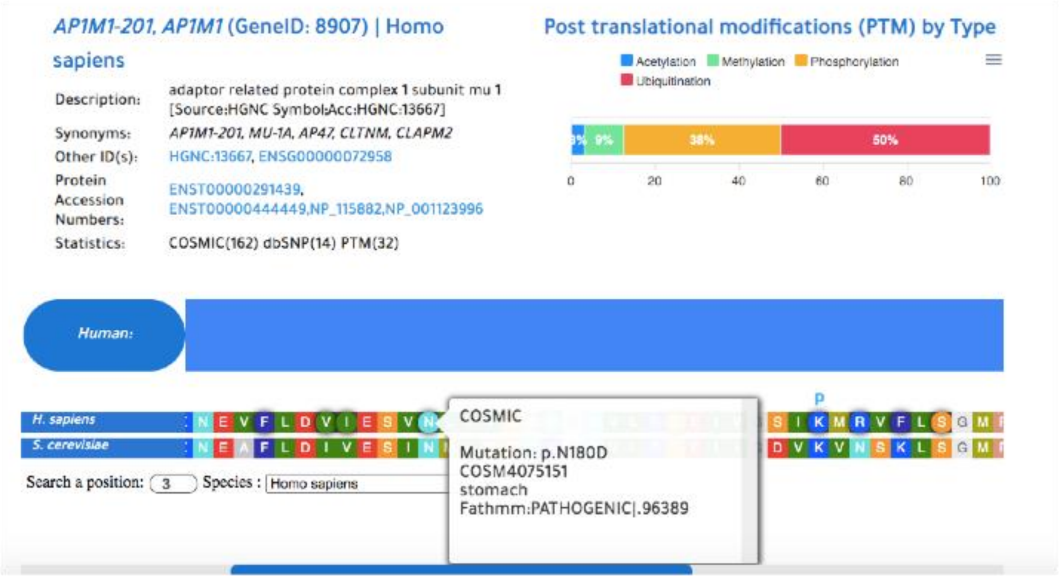


**Figure S2:** Amino sequences of Apm1p from *Saccharomyces cerevisiae* (NP_015064.1) were retrieved from NCBI and were then submitted to ConVarT for Needleman-Wunsch global alignment**.** ConVarT has successfully discovered the closest human gene (human AP1M1, the human ortholog of *S. cerevisiae*

Apm1p), displaying amino acid variants from humans on the pairwise alignment of human AP1M1 and *S. cerevisiae* Apm1p. Shown is the p.N180D variant from COSMIC between human AP1M1 and yeast Apm1p.

**The entry of new variant and phenotypic data**

ConVarT is an open-access platform, enabling scientists to easily submit variant and/or phenotype data to ConVarT to disseminate the knowledge they produce. To submit a new variant and/or phenotypic data, they need to visit https://convart.org/pages/Submit.php and submit the relevant data as displayed on the website. They can submit the reference paper if available, thus we can include the reference link to visitors. If no published reference is available at the time of submission, they can still submit it to ConVarT. Their submission will be cited as personal communication. Once they submit the data, we manually review the submission. Once they are approved, they will be automatically integrated into ConVarT.

| Identifier Name | Description |
|---|---|
| NCBI Gene ID | NCBI Gene ID (e.g. 6301) |
| Gene Symbol | Current Gene Symbol (e.g. *SARS*, *sars-1*) |
| Gene Synonyms | Previous Gene Symbols (e.g. *SERRS*) |
| HGNC | HUGO Gene Nomenclature Committee (e.g. HGNC:10537) |
| ENSEMBL GENE ID | ENSEMBL Gene IDs for any specie in our database (e.g. ENSG00000031698, ENSMUSG00000068739, ENSRNOG00000020255, |

| | |
|---|---|
| | ENSDARG00000008237, ENSPTRG00000001043, ENSMMUG00000021837) |
| Variant ID | Reference SNP ID (e.g. rs1553178049) |
| Protein Acc. Number | NCBI Protein Accession Numbers (e.g. NP_006504.2, XP_006233198.1) |
| MGI ID | MGI-Mouse Genome Informatics (e.g. MGI:102809) |
| ZFIN ID | ZFIN The Zebrafish Information Network (e.g. ZDB-GENE-040831-1) |
| FB Gene ID | FlyBase Gene ID (e.g. FBgn0031497) |
| WB Gene ID | WormBase Gene ID (e.g. WBGene00005663) |

**Supplementary Table 17:** Shown table is the list of the identifiers that can be used as input for gene searching on ConVarT.

**Supplementary Tables**

Table 1: Human and Mouse with Matching Variants (All)
Table 2: Human variants and *C. elegans* Matching Variants (All)
Table 3: *C. elegans* Variants Without Human Equivalent Variants (SIFT and Disease added)
Table 4: *C. elegans* Phenotypic Variants without Human Equivalent Variants (SIFT and Disease added)
Table 5:Human Benign Variants and *C. elegans* Phenotypic Variants
Table 6:Human Benign Variants (ClinVar) and C. elegans Unknown Variants
Table 7:Human Benign Variants (ClinVar) and Mouse Phenotypic Variants
Table 8:Human Benign Variants (ClinVar) and Mouse Unknown Variants
Table 9:Human Variants and *C. elegans* Matching Variants (No mouse orthologous genes)
Table 10:Human Variants and Mouse Double Matching Variants (No *C. elegans* Matching Variants)

Table 11:Human Pathogenic variants and *C. elegans* Phenotypic variants
Table 12:Human Pathogenic variants and *C. elegans* Unknown variants
Table 13:Human Pathogenic variants and Mouse Phenotypic variants
Table 14:Human Pathogenic variants and  Mouse Unknown variants
Table 15: Mouse Variants Without Human Equivalent Variants (SIFT and Disease added)
Table 16:Mouse Phenotypic Variants  Without Human Equivalent Variants (SIFT and Disease added)

### Material and Methods

#### Homologs and Orthologs of Human Genes and Sequence Retrieval

For ConVarT, we first created a gene homology list for the following organisms: *Pan troglodytes* (Chimps), *Macaca mulatta* (Macaca), *Mus musculus* (Mouse), *Rattus rattus* (Rat), *Xenopus laevis* (Xenopus), *Danio rerio* (Zebrafish), *Drosophila melanogaster* (Fruitfly) and *Caenorhabditis elegans* (Worm). We used several resources to compile a gene homology list. We obtained the gene homology list from the Mouse Genome Informatics (MGI) (http://www.informatics.jax.org/downloads/reports/HOM_AllOrganism.rpt) for Chimps, Macaca, Mouse, Rat and Xenopus. However, we noticed that this homology list is missing counterparts of human genes for Chimp and Macaca. In a brief, to find the homologs of missing genes for Chimp and Macaca, we locally conducted BLAST analysis with the default BLAST protein parameters (blastp) version 2.2.28+ (1). For this, we downloaded the protein sequence as a FASTA file for humans and the complete genomic sequences of Chimps and Macaca from NCBI and used human FASTA files as input to locally run BLAST analysis against the Chimps and Macaca genomes. We downloaded the human-Zebrafish homology genes list from the Zebrafish Information Network (ZFIN) (Here is the website for the download https://zfin.org/downloads/human_orthos.txt ) (2). We retrieved the human - Drosophila homology matching list from DRSC Integrative Ortholog Prediction Tool (DIOPT) on 1st March 2019 (the website for the download https://www.flyrnai.org/cgi-bin/DRSC_orthologs.pl ) (3). We excluded the human and Drosophila homolog pairs with low scores (score < 2). Finally, we created a human-*C. elegans* orthology list by using the in-house protein BLAST (blastp) pipeline. We performed reciprocal BLAST analysis for *C. elegans* and humans. First, *C. elegans* protein sequences were obtained from UniProt and were used as input against the human complete human genome. We performed a similar BLAST analysis for human proteins as input against the human complete *C. elegans* genome. If genes received the highest-scoring in the reciprocal blast analysis, we accepted them as orthologs. If the gene did not receive the highest-scoring in one of these two blast analysis, we took an alternative way. Non-matching genes from the BLAST analysis (*C. elegans* proteins as input) were compared with the list of *C. elegans*-human orthologs from the human OrthoList 2 (OL2) (4). If our BLAST result matches with the OrthoList 2, we consider them as orthologs.  In a case of no matching, we manually searched for them in the literature and integrated them into the list of *C. elegans*-human orthologs.

**Retrieval of Genetic Variants for Humans, Mice and *C. elegans* and Posttranslational modification (PTMs)**

We retrieved the ClinVar variant data from ftp://ftp.ncbi.nlm.nih.gov/pub/clinvar/ using the in-house Python3 script (https://github.com/thekaplanlab/ConVarT_pipeline ) (5) and we regularly downloaded the ClinVar databases. The last update was August, 2021. There are no variant IDs available for 765 records and they will appear "N/A" on the ConVarT website. We downloaded the genome Aggregation Database (gnomAD) v2.03 (Here is the website for the download: http://gnomad.broadinstitute.org/) it is already in GRCh37), COSMIC (Here is the website for the download: https://cancer.sanger.ac.uk/download we advertently select GRCh37), dbSNP (Here is the website for the download: https://ftp.ncbi.nih.gov/snp/ and PhosphoSitePlus (IDK) (Here is the website for the download: https://www.phosphosite.org/Supplemental_Files) (6–9). The details can be found on https://github.com/thekaplanlab/ConVarT_pipeline. Variants together with phenotypic data for Mouse and *C. elegans* were generously shared by Mutagenetix (Here is the website for the download https://mutagenetix.utsouthwestern.edu/about.cfm) and Wormbase (10, 11). We retrieved mouse phenotypic data together with corresponding variants from the website of Australian Phenomics (the website for download https://pb.apf.edu.au/phenbank/homePage.html ). Our ConVarT database currently presents over 40,000,000 genetic variations.

**Codes Availability, Data Sharing and ConVarT website**

We published three repositories, Convart_Pipeline (https://github.com/thekaplanlab/Convart_Pipeline ), ConVarT_Web https://github.com/thekaplanlab/ConVarT_Web ) and ConVarT_Analysis (https://github.com/thekaplanlab/ConVarT_Analysis). The former repository can be used to reproduce our database, while the latter consists of the source code for the ConVarT website. In ConVarT_Analysis, we provided codes used to generate figures and statistical analysis. Data sharing is a key to advance science and increase reproducibility in Science. We share all data and resources at the following website http://convart.org/pages/Downloads.php .

| Databases | Number of Variants | Download Date | References |
|---|---|---|---|
| gnomAD | 17,009,216 | August, 2021 | (9) |

| | | | |
|---|---|---|---|
| ClinVar | 2,192,865 | August, 2021 | (5) |
| COSMIC | 12,497,561 | September, 2021 | (8) |
| dbSNP | 1,086,546 | August, 2021 | (6) |
| Wormbase | 252,260 | August, 2021 | (11) |
| Mutagenetix | 481,050 | August, 2021 | (10) |
| TOPMed | 25,275,907 | August, 2021 | (20) |
| PhosphoSitePlus | 383k | June 29, 2019 | (7) |
| Phenomics Mice | 50,800 | August, 2021 | The Australian Phenome Bank * |

**Table 18**: * We downloaded the mouse variants and phenotypic data from https://pb.apf.edu.au/phenbank/homePage.html

**Retrieval of Protein Sequences for Humans, Mice and *C. elegans* and Multiple Sequence Alignments**

We used an in-house script to retrieve human transcripts annotated in ClinVar from the GPFF file of the human genome with version GRCh37. Toward this goal, “NM” identifiers are matched to “NP” or “XP” protein accession numbers one-to-one using an “NM to NP/XP” list. With these protein accession numbers, we next used an in-house script to retrieve the correct transcript of protein isoforms from the human reference protein sequence FASTA file. For gnomAD and COSMIC databases, the protein sequences of variants were retrieved ENST accession numbers from Ensembl with the same in-house python script. Duplicated accession numbers from gnomAD and COSMIC were discarded. We downloaded the GPFF file of the chimp, Macaca, rat, mouse, zebrafish, *C. elegans, D. melanogaster,* and frog genomes and retrieved the amino acid sequences of human orthologous genes together with protein accession numbers. We created small files storing the protein sequences of homolog genes from human, chimp, Macaca, rat, mouse, zebrafish, *C. elegans, D. melanogaster,* and frog, followed by comprehensive multiple sequence alignments (MSAs) of these protein sequences with ClustalW v2.1 pairwise alignment. All created sequences are processed in Python3 and stored in MySQL (12). Our database currently hosts over 2,379,397 different combinations of MSAs.

#### Curation of Transcripts

We retrieved the correct human protein transcript or protein isoform for enabling ConVarT to visualize the variations into corresponding amino acid positions, it is needed to use protein identifiers that are annotated in the variant cataloging databases and post-translational modification (PTMs) from PhosphoSitePlus data set.

#### Generation of Matching Variants

We focus on equivalent variants namely “matching variants” (MatchVars). We first define matching variant terms as two or more aligned variants sharing the same reference amino acid and potentially similar functional impact (phenotypic, pathogenic, and benign). The potential functional impact of variants were determined according to BLOSUM similarity matrix. Amino acid changes between similar amino acids were considered as conservative changes, whereas the changes between dissimilar amino acids were treated as radical changes. There are a number of rules for a variant to be included as an MatchVar. 1) Upon multiple sequence alignments or pairwise sequence alignment, amino acid residues in the corresponding position between two orthologous genes from different organisms must be the same. 2) Both the amino acid residues in the corresponding position should undergo the same type of change. Both changes should either be conservative or radical for both variants to be considered as orthologous variants (13). Unless they are in accordance with the BLOSUM similarity matrix, amino acid-changing variants have not been considered as MatchVar. If both variants have the same reference amino acids and have the same type of amino acid changes (conservative or radical), they are considered as orthologous variants. For example, R897Q substitution in human ERBB2 encoding protein NP_004439 is aligned with R898C variation in mouse Erbb2 encoding protein NP_001003817 in multiple sequence alignment. Although both variants have the same reference amino acid, while the substitution of arginine to glutamine is considered as conserved, conversion of arginine to cysteine is a more radical change. Therefore, these were not considered as MatchVar in our analysis.

To find all matching variants between human and mouse and human and *C. elegans*, we used multiple sequence alignments (MSAs) data and variants from human (ClinVar, dbSNP, COSMIC and gnomAD), mouse (Mutagenetix and Australian Phenomics Bank) and *C. elegans* (Wormbase). We first determined if the reference amino acids at the corresponding positions are the same. Then we compared the amino acid changing variants at the corresponding positions from humans with those from mouse and *C. elegans* by using an amino acid substitution conservation table created based on BLOSUM similarity matrix (13). If both variants have the same reference amino acids and have the same type of amino acid changes (conservative or radical), they are considered as orthologous variants. Supplementary tables presenting orthologous variants include all possible protein alignments.

**Analysis of Variants and Statistical Analysis**

Since ConVarT focuses on amino acid variants, we included only missense and nonsense variant data, excluding all other types of variants including a change from the stop codon to an amino acid to the analysis. We also excluded variants with no Refseq protein ID. We included variants if at least one of protein IDs, amino acid change, amino acid position, or clinical significance is unique. Others were regarded as duplicates and excluded from further analysis. We discarded variants with more than one amino acid change. For *C. elegans* variant data, we changed "opal stop", "amper stop" and "ochre stop" terms to the asterisk (*) for consistency. For all variants, we changed three-letter amino acid codes to one letter codes (for example, Ala>A). We excluded the data including any string other than amino acid single letter or three letter codes in the amino acid change columns. We converted the variants annotated with Ensembl transcript ID and Ensembl protein ID to RefSeq protein ID. For data with Ensembl transcript ID, we first converted them to Ensembl protein ID via BioMart (14). Then we matched Ensembl protein IDs to RefSeq IDs through protein sequences. We retrieved protein sequences from Ensembl and NCBI FTP sites. We converted CCDS IDs in mouse variant data downloaded from the Australian Phenome Bank to RefSeq ID by matching protein sequences. We downloaded CCDS-protein sequence annotation data from the NCBI FTP website.

**Calculation of SIFT Scores**

To calculate SIFT scores of *C. elegans* variants, we first generated a VCF file, which provides the genomic location of variants, from protein IDs, amino acid change, and position information using Variant Effect Predictor (VEP) tool from Ensembl (https://www.ensembl.org/Tools/VEP) (15, 16). Then, we used the SIFT 4G annotator application to find SIFT scores by using the VCF file as input. For variants taken from Mutagenetix, we used the Ensembl VEP tool for SIFT scores. Other prediction scores for mouse variants, including Polyphen scores and SIFT scores for APF mouse mutations were available in the corresponding variant data. To generate tables for mouse and *C. elegans* variants having no corresponding variant in humans, we excluded orthologous variants from variant lists. For ClinVar variants, allele frequency data from various sources including gnomAD, TOPMed, and ExAC were retrieved from ClinVar and integrated to ClinVar variant data. All other analyses were performed using R and the source code is available on GitHub (https://github.com/thekaplanlab/ConVarT_Analysis).

**Needleman-Wunsch global alignment for the visualization of human variants**

Scientists can perform homology searches on the ConVarT to visualize the distribution of human variants in the protein sequences they submit (**Figure S3)**. ConVarT was designed to accept the sequence as a FASTA (https://convart.org/ ) in the search box. Once they submit the amino acid sequence, the ConVarT will perform the Needleman-Wunsch global alignment to find the closest human gene, and will display human variants on the pairwise sequence alignment between the submitted protein sequence and the closest amino acid sequence from humans (17). The use of CRISPR has exponentially increased, and there is a need to easily inspect the presence of human variants on the gene of their interest. We therefore believe this feature will be useful to many who may wish to find human variant distributions on the gene of their interest.

**Human Protein Domains**

With protein accession numbers from ClinVar, gnomAD, dbSNP, and COSMIC, we curated human protein transcripts from NCBI GRCh37 and Ensembl protein sequence FASTA file ftp://ftp.ebi.ac.uk/pub/databases/Pfam/Tools/ folder. We used them as a query file in the Pfam-scan script when they were downloaded. We performed Pfam script with default parameters to obtain Pfam domains, family and clan information, Pfam profiles and HMM models from ftp://ftp.ebi.ac.uk/pub/databases/Pfam/current_release link with a version of Pfam 32.0. We obtained a total of 68,006 Pfam domains total with 6,433 unique Pfam IDs for 25,484 unique transcript IDs. We set 1e-01 e-value as a threshold and only display those protein domains on ConVarT.

**Use of Tables and Other Components on the ConVarT Website**

ConVarT uses a specialized and sortable table plug-in for jQuery to show the large variant and PTMs data set. Here, also, users can reach the original record or variant id using the links that are shown as blue to access and display the record on the specific database for detailed information.

**Sequence Viewer**

ConVarT displays multiple sequence alignments that are created in advance to reduce search time. The user can search genes across eight non-human species. However, MatchVar focuses on only human, mouse and *C. elegans*. In that regard, ConVarT has specialized sequence alignment and domain viewer to make the MSA results and accessing of variations in the corresponding position of amino acids much more user-friendly and ready-to-use. Therefore, that component is based on JavaScript and jQuery with HTML5 and CSS3 technologies. In addition to these points, owing to the need to search a certain amino acid position in any species transcript, the small text box enables researchers to directly find out the amino acid or variation that they are interested in. The coloring of the amino acids is based on RasMol Shapely Colors.

**Interactive Graphs**

We integrated interactive stacked bar charts into ConVarT to visualize the fractions of variants from ClinVar and others or post-translational modifications (PTMs) per gene. For this purpose, ApexChart, an open-source JavaScript library, was used to display the variations or PTMs stored on MySql with help of PHP. Users can download the high quality of the interactive graphs using the menu button at the right corner of the chart.

**Disease Genes (DisGeNet) Analysis and Displaying on the web**

Search with disease names was integrated into ConVarT. Toward this goal, we downloaded disease-associated genes in 26 different disease categories curated by DisGeNet. Here is the link for the download: http://www.disgenet.org/static/disgenet_ap1/files/downloads/curated_gene_disease_associations.tsv.gz (18). We used the in-house Python3 script to match disease genes with corresponding orthologous genes from 8 non-human species, thus making it easy to scan for genes with disease names. For example, searching with "Joubert syndrome " will show up all Joubert-syndrome associated genes, thus users can click on the gene of interest. We created a pie chart showing the distributions of all the homologs of human disease-associated genes in 8 non-human species across 26 disease categories using Python3, matplotlib, and pandas packages. Besides, the nomenclature of disease categories is retrieved from MeSH Browser under https://meshb.nlm.nih.gov/treeView link. On the web page of a gene, ConVarT enumerates the diseases that are associated with the searched genes. In the case of displaying a gene from different-organisms, ConVarT displays the lists of diseases of the human homolog of the searched gene.

**Comparative visualization of variants between human and other organisms**

We used the trackViewer, a Bioconductor package, to visualize protein domains along with variants in **Figure 3** (19). The codes used in the production of **Figure 3** can be found on the following site https://github.com/thekaplanlab/ConVarT_Analysis .

**Addition of AlphaFold**

With the availability of the accurate protein structure prediction database AlphaFold for humans, mice, and *C. elegans*, the AlphaFold database was added as a link for each gene. Users can visit the database to compare protein structures of orthologous genes in these organisms (21).

*Nat. Genet.*, **3**, 266–272.
