## Supplementary figures and images for "ConVarT: a search engine for matching human genetic variants with variants from non-human species"

### Supplementary Figure 1

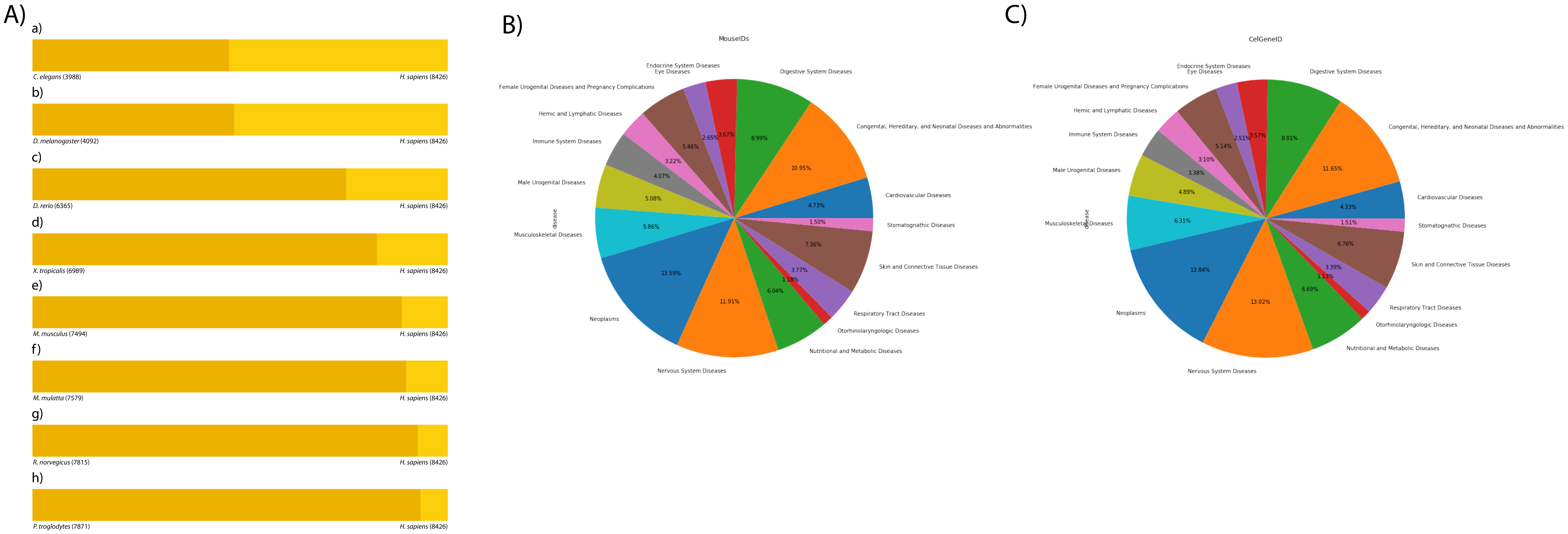

### Supplementary Figure 2

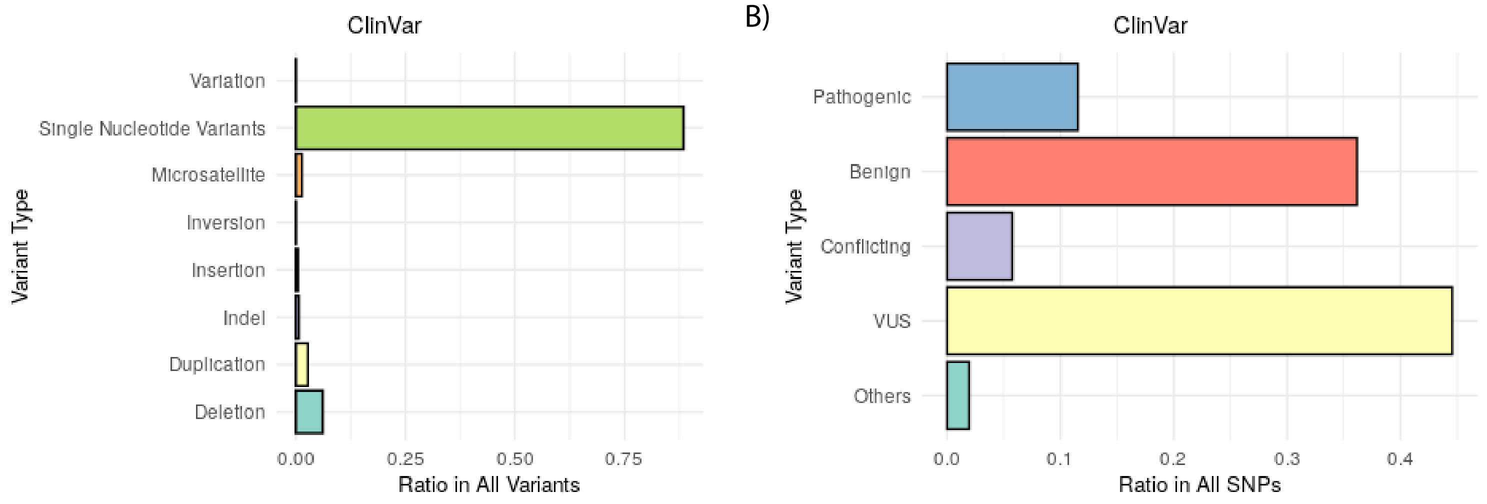
