## Supplementary Figure 3 for "ConVarT: a search engine for matching human genetic variants with variants from non-human species"

### *AP1M1-201, AP1M1* (GeneID: 8907) | Homo sapiens

| | |
|---|---|
| Description: | adaptor related protein complex 1 subunit mu 1 [Source:HGNC Symbol;Acc:HGNC:13667] |
| Synonyms: | *AP1M1-201, MU-1A, AP47, CLTNM, CLAPM2* |
| Other ID(s): | HGNC:13667, ENSG00000072958 |
| Protein Accession Numbers: | ENST00000291439, ENST00000444449,NP_115882,NP_001123996 |
| Statistics: | COSMIC(162) dbSNP(14) PTM(32) |

### Post translational modifications (PTM) by Type

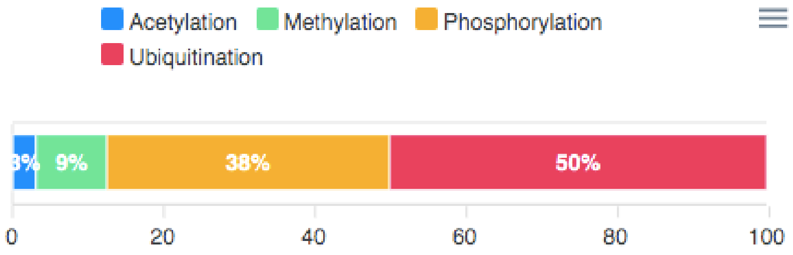


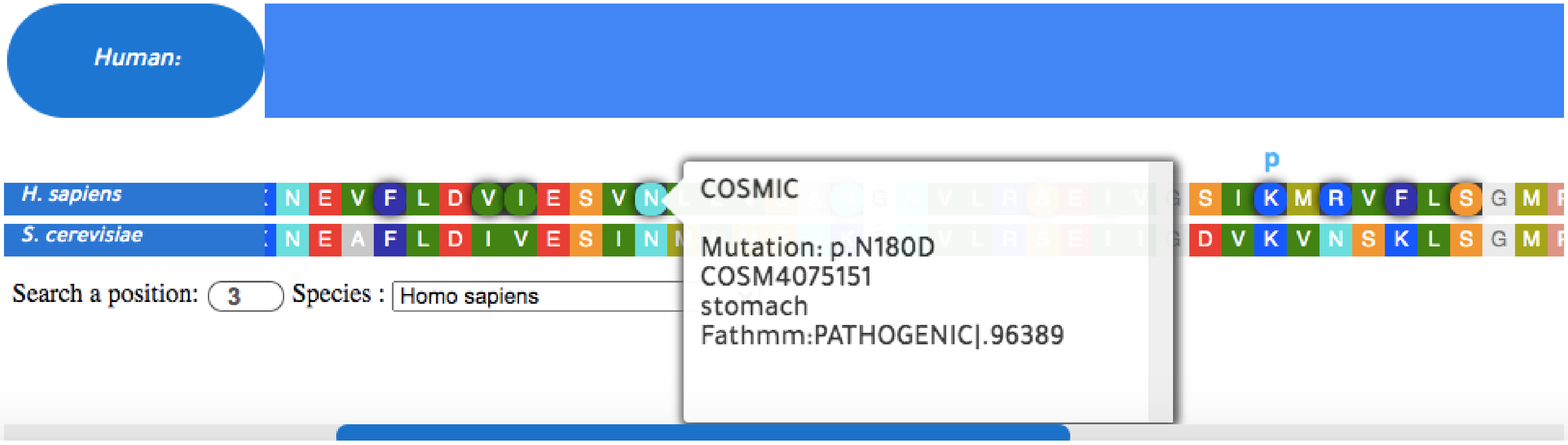
